# HD-PTP/PTPN23 hypomorphic mice display lipodystrophy

**DOI:** 10.1101/2022.08.02.502401

**Authors:** Brian A. Davies, Johanna A. Payne, Cole P. Martin, Destiny Schultz, Bennett G. Childs, Cheng Zhang, Karthik Jeganathan, Ines Sturmlechner, Thomas A. White, Alain de Bruin, Huiqin Chen, Michael A. Davies, Sarah Jachim, Nathan K. LeBrasseur, Robert C. Piper, Hu Li, Darren J. Baker, Jan van Deursen, David J. Katzmann

**Affiliations:** Department of Biochemistry and Molecular Biology; Department of Pediatric and Adolescent Medicine; Department of Molecular Pharmacology and Experimental Therapeutics; Robert and Arlene Kogod Center on Aging, Mayo Clinic, Rochester, Minnesota, 55905, United States; Department of Pediatrics, University of Groningen, University Medical Center Groningen, Groningen, 9713 AV, The Netherlands; Department of Pathobiology, Faculty of Veterinary Medicine, Utrecht University, Utrecht, 3584 CL, The Netherlands; Department of Biostatistics, Division of Quantitative Sciences; Department of Melanoma Medical Oncology, Division of Cancer Medicine, University of Texas M. D. Anderson Cancer Center, Houston, Texas, 77030, United States; Department of Molecular Physiology and Biophysics, University of Iowa, Iowa City, Iowa, 52242, United States

## Abstract

Endosomal Sorting Complexes Required for Transport (ESCRTs) drive reverse topology membrane remodeling events including the formation of intralumenal vesicles within multivesicular bodies, the budding of retroviruses from the plasma membrane, and the scission of the cytokinetic bridge. It has been difficult to study the physiological relevance of this machinery in mammals because many contributing components are essential for viability. To bypass this problem we used combinations of knockout (−), hypomorphic (H) and wildtype (+) alleles to generate a series of mice with a gradual reduction of HD-PTP (product of *PTPN23*), an ESCRT-associated protein known to cause embryonic lethality when fully depleted. Whereas *PTPN23^-/H^* mice died shortly after birth, *PTPN23*^H/H^ mice developed into adulthood but had reduced size, lipodystrophy, and shortened lifespan. Analysis of 14-day inguinal adipose tissue indicated reduced expression of adipogenesis markers, and *PTPN23* knockout preadipocytes similarly display reduced adipogenesis *in vitro.* Defects in insulin-stimulated signaling were apparent in differentiated *PTPN23* knockout adipocytes and *PTPN23^H/H^* inguinal adipose tissue *in vitro*, correlating with reduced levels of insulin signaling hallmarks observed in adult *PTPN23^H/H^* inguinal adipose tissue *in vivo.* Whereas the ESCRT machinery have been suggested to downregulate signaling, these results indicate that HD-PTP promotes insulin-induced signaling in, as well as differentiation of, inguinal adipose tissue. These results revealed unexpected roles for HD-PTP in promoting fat accumulation in mammalian cells through supporting insulin signaling, adipogenesis, and lipid droplet formation.

## Introduction

The Endosomal Sorting Complexes Required for Transport (ESCRTs: ESCRT-0, I, II and III) facilitate reverse topology membrane remodeling events that bud vesicles away from the cytoplasm, facilitate membrane scission from within a membrane tubule, or accomplish membrane repair (reviewed [1–8]). The biological processes facilitated by ESCRTs include intralumenal vesicle formation during multivesicular body biogenesis and exosome biogenesis, abscission of the cytokinetic bridge, closure of the autophagosome, repair of membrane lesions, and maintenance of nuclear envelope integrity. Consistent with roles in diverse, important cellular processes, disrupted ESCRT function through knockout of ESCRT-0 (*hrs*) [9], ESCRT-I (*tsg101*) [10], ESCRT-III (*chmp5*) [11] or the ESCRT-associated factor HD-PTP (*ptpn23*) [12] result in embryonic lethality in mouse models. Moreover, mutations altering ESCRT function have been linked to congenital disorders including spastic paraplegia (SP80, *UBAP1;* SP53, *Vps37A*) [13–15], childhood cataracts (CTRCT31, *CHMP4B*) [16, 17], frontotemporal dementia (FTD/ALS17, *CHMP2B*) [18–21], pontocerebellar hypoplasia (PCH8, *CHMP1A*) [22], cerebellar hypoplasia, cataracts, impaired intellectual development, congenital microcephaly, dystonia, dyserythropoietic anemia, and growth retardation (CIMDAG, *VPS4A*) [23–25] and neurodevelopmental disorder with structural brain anomalies, seizures and spasticity (NEDBASS, *HDPTP/PTPN23*) [26–30] (also reviewed in [31]). While the ESCRTs serve important roles facilitating development and maintaining organismal homeostasis, the lack of a viable ESCRT-deficient mouse model has precluded a deeper understanding of how this is achieved.

The Bro1 Domain family members - including HD-PTP [32–37] and ALIX [38–41] in mammalian systems and Bro1 in yeast [42] – are ESCRT-associated factors facilitating ESCRT-mediated processes. These factors contribute to cargo recognition through 1) binding cargo directly [38, 40, 43, 44], 2) binding ubiquitin coupled to cargo [45–48], and 3) binding ESCRT-0 and ESCRT-I [34, 38, 43, 47, 49–54], factors that contribute to cargo recognition. These Bro1 Domain family members also function later during ESCRT-mediated processes through 1) promoting ESCRT-III assembly via activating CHMP4/Snf7 [40, 55, 56], 2) recruiting ubiquitin isopeptidases to recycle ubiquitin from cargo [52, 57–60], and 3) facilitating ESCRT-III-driven membrane remodeling by activating the Vps4 ATPase [48, 61]. These varied activities position Bro1 Domain family members as critical factors regulating ESCRT-driven processes [62]. HD-PTP and ALIX mediate distinct ESCRT processes. Specifically, ALIX has been implicated in ESCRT-mediated viral budding (e.g., surface budding of HIV) [38, 40], cytokinesis [63, 64], exosome biogenesis [65–67], and plasma membrane repair [68]. *PDCD6IP* (ALIX) knockout mice models display altered brain development but are viable and display normal lifespan [69, 70]. In contrast, a *PTPN23* (HD-PTP) knockout mice model exhibits embryonic lethality [12]. Moreover, *PTPN23*^-/+^ mice display increased tumorigenesis [71], and HD-PTP has been implicated in MVB sorting for lysosomal degradation [33, 49] as well as in signaling of Toll receptors [72], cilia-associated signaling [73], SMN complex function [74], and neuron guidance and pruning [75]. These differential phenotypes of *PTPN23* and *PDCD6IP* knockout mice suggest that HD-PTP contributes to ESCRT-mediated processes required for survival while ALIX does not.

Better understanding HD-PTP functions contributing to organismal homeostasis is precluded by the embryonic lethality of *PTPN23* knockout animals [12]. To gain further insights into the physiological function of HD-PTP in mammalian systems, we created a hypomorphic allele of *PTPN23* (H). *PTPN23^H/H^* animals are viable but exhibit reduced survival and lipodystrophy. Mechanistically, HD-PTP deficiency contributes to lipodystrophy through both cell extrinsic and cell autonomous mechanisms, including altered insulin signaling. These findings uncover an unexpected role for HD-PTP in organismal homeostasis by impacting fat accumulation.

## Results

### Generation of PTPN23 allelic series

Homozygous deletion of *PTPN23* confers embryonic lethality at stage e9 in mice [12]. To circumvent this lethality and assess the contribution of HD-PTP to organismal physiology, an allelic series of animals with varied gene dosage was generated and assessed for the minimum level of HD-PTP required for viability. A hypomorphic allele (*PTPN23^H^*) was generated by insertion of a neomycin resistance cassette containing cryptic splice acceptor and donor sequences between exons 6 and 7 (Figure 1A) [76]. When spliced into the mRNA, the neomycin resistance cassette contains a premature stop codon that will truncate HD-PTP within the Bro1 Domain. Non-cryptic mRNA processing bypasses the neomycin resistance cassette, leaving otherwise normal mRNA. With the neomycin splice being non-obligatory, splicing from exon 6 to 7 bypassing this exon produces lower numbers of normal mRNA transcripts, thereby reducing overall protein expression. To allow for removal of this hypomorphic insert, FRT sites were introduced flanking the neomycin resistance cassette. Flp-mediated recombination reverts the hypomorphic allele and generates an inducible knockout allele (*PTPN23^flox^*). CRE-mediated recombination of LoxP sites excises exons 5 and 6 generating a functionally null allele (*PTPN23^-^*). Heterozygous animals were crossed and surviving progeny genotyped (Figure 1B). Normal Mendelian patterns were observed in the context of *PTPN23^H/+^* intercrosses and *PTPN23^H/+^* crossed to *PTPN23^-/+^*, while no *PTPN23^-/-^* animals were recovered from *PTPN23^-/+^*intercrosses, which was expected with previous studies [12]. These results indicate that the minimal gene dosage sufficient to support embryogenesis is *PTPN23^H/−^*. However, *PTPN23^H/-^* animals did not survive beyond one day postnatally (data not shown). We found that the minimal gene dosage for physiological studies is the homozygous hypomorphic allele animal *(PTPN23^H/H^*). While viable into adulthood, *PTPN23^H/H^* mice were infertile, which required maintaining the colony through *PTPN23^H/+^* intercrosses. *PTPN23^H/H^* and *PTPN23^+/+^* offspring were analyzed for comparisons. As expected, analysis of HD-PTP expression in newborn *PTPN23^H/H^* animals revealed reductions in HD-PTP protein levels. HD-PTP expression in the brain was reduced 63% in *PTPN23^H/H^* neonates compared to 42% and 77% reductions in *PTPN23^-/+^* and *PTPN23^H/-^* pups, respectively (Figure 1C, Figure S1). Further inspection of HD-PTP expression in *PTPN23^H/H^* neonates revealed reductions in HD-PTP in all tissues examined (Figure 1D) with many tissues exhibiting more than 50% reductions. Survival of male and female *PTPN23^H/H^* mice was significantly decreased from *PTPN23^H/+^, PTPN23^-/+^*, and *PTPN23^+/+^* male mice (Figure 1E); there were no deaths in the latter three groups for at least one year. *PTPN23^flox/flox^* and *PTPN23^+/+^* mice were indistinguishable from each other, indicating the hypomorphic allele could be reverted to complement mutant phenotypes. While heterozygous *PTPN23^-/+^* animals have been reported to display increased spontaneous mesenteric lymphoma and lung adenoma incidence by 72 weeks of age [71], we found no cancer predisposition in our *PTPN23^-/+^, PTPN23^H/+^*, or *PTPN23^H/H^* mice (data not shown). The primary cause of death for *PTPN23^H/H^* animals is unclear as a variety of conditions – including unexplained decreased mass - led to termination through IAUCUC protocol guidelines. In total, these analyses indicate that HD-PTP function is required for embryogenesis as well as maintenance of normal physiology in the adult mouse.

**Figure 1.**
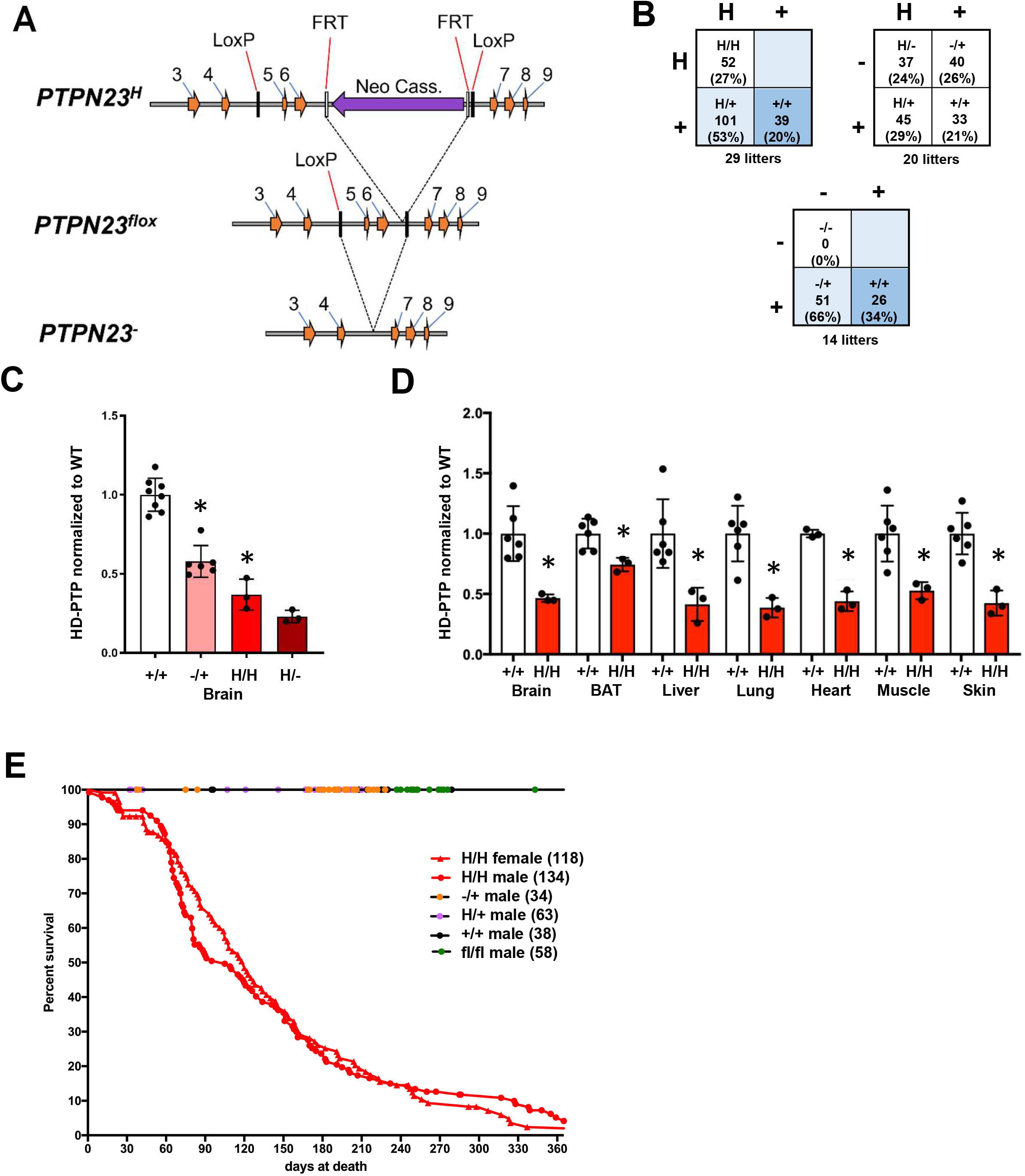
Generation of *PTPN23* hypomorphic allele. **A.** Schematic representation of the gene targeting strategy employed to generate hypomorph (**H**), conditional (**flox**) and knockout (**-**) alleles of *PTPN23*. Insertion of exogenous allele with moderate splice acceptors between exons 6 and 7 reduces but does not eliminate HD-PTP expression. Flp recombinase expression drives excision of the exogenous exon via FRT sites but leaves a LoxP site between exon 6 and 7. A second LoxP site was introduced between exons 4 and 5 in the hypomorphic allele. Cre recombinase expression results in excision of exons 5 and 6 via LoxP sites generating a null allele by altering the open reading frame. **B.** Progeny of crosses between *PTPN23^H/+^* animals, *PTPN23^H/+^* and *PTPN23^-/+^* animals, and *PTPN23^-/+^* animals. Crosses of *PTPN23^-/+^* animals did not produce *PTPN23^-/-^* progeny. Crosses of *PTPN23^H/+^* animals or *PTPN23^H/+^* and *PTPN23^-/+^* animals yield progeny with expected mendelian distributions; however, *PTPN23^H/-^* pups failed to survive beyond the first day. **C.** Analysis of HD-PTP expression in brain tissue from *PTPN23^+/+^, PTPN23^-/+^, PTPN23^H/H^* and *PTPN23^H/-^* newborn pups. HD-PTP expression is significantly reduced in *PTPN23^-/+^* compared to *PTPN23^+/+^*, and *PTPN23^H/H^* compared to *PTPN23^-/+^*. * indicates two-tailed t-test p-value ≤ 0.05. **D.** Analysis of HD-PTP expression in *PTPN23^+/+^* and *PTPN23^H/H^* tissues from newborn pups: brain, brown adipose tissue (BAT), liver, lung, heart, muscle, and skin. HD-PTP expression is significantly reduced in all *PTPN23^H/H^* tissues examined. * indicates two-tailed t-test p-value ≤ 0.05. **E.** Survival curves of female and male *PTPN23^H/H^* animals compared to male animals of genotypes *PTPN23^-/+^*, *PTPN23^H/+^, PTPN23^+/+^* and *PTPN23^flox/flox^*. Both male and female *PTPN23^H/H^* animals exhibit reduced survival compared to *PTPN23^+/+^* (Log Rank Test p value < 0.0001).

### PTPN23^H/H^ mice exhibit lipodystrophy

In addition to decreased survival, *PTPN23^H/H^* mice have reduced size and mass throughout their lives (Figure 2A-C). This phenotype is not observed in *PTPN23^flox/flox^* mice (Figure S2), demonstrating linkage to the hypomorphic allele. To further explore the reduced mass phenotype in *PTPN23^H/H^* mice, body composition was assessed using Echo-MRI in animals at 7-10 weeks of age. This analysis revealed significant reductions in body fat composition in both female and male *PTPN23^H/H^* mice (Figure 2D). Comprehensive Laboratory Animal Monitoring system (CLAMs) was employed to further assess behavioral or metabolic changes that might contribute to this altered composition. While female *PTPN23^H/H^* mice have reduced adjusted food intake and increased activity compared to *PTPN23^+/+^* mice, similar differences were not observed in male mice (Figure 2E and 2F). Thus, reduced food consumption or increased activity seemingly are not the primary drivers of altered body composition in *PTPN23^H/H^* mice. Both female and male *PTPN23^H/H^* mice exhibit increased energy expenditure when normalized to total body mass compared to *PTPN23^+/+^* controls (Figure 2G). This difference may result from altered body composition as normalizing energy expenditure to lean body mass rather than total body mass eliminated this effect in male mice (Figure 2H). These analyses did not identify an explanation for the reduced fat content of the *PTPN23^H/H^* mice, which warranted further examination of the adipose tissue.

**Figure 2.**
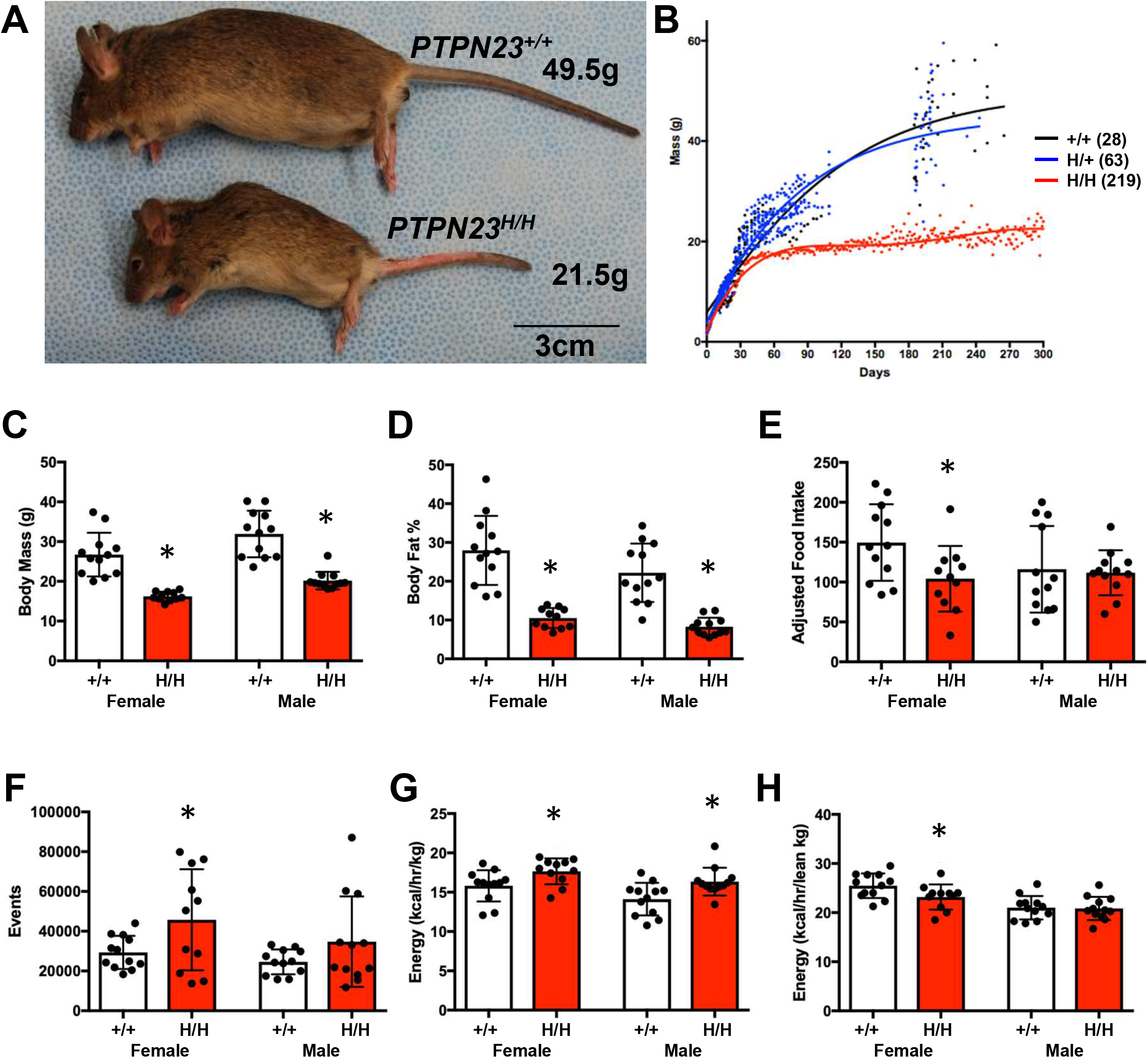
*PTPN23^H/H^* animals exhibit reduced mass and fat composition. **A.** Representative image of adult *PTPN23^+/+^* and *PTPN23^H/H^* male mice. **B.** Mass in *PTPN23^+/+^*, *PTPN23^H/+^* and *PTPN23^H/H^* male animals over first year. **C, D.** Total body mass (**C**) and total body fat composition analysis by ECHO MRI (**D**) of 6-week old *PTPN23^+/+^* and *PTPN23^H/H^* adult male and female animals. **E-H.** Adjusted food intake (**E**), total activity (**F**), and total energy expenditure normalized to total body mass (**G**) or lean body mass (**H**) were assessed for 6-week old *PTPN23^+/+^* and *PTPN23^H/H^* male and female animals. * indicates two-tailed t-test p-value ≤ 0.05.

Analyses of fat deposits in adult animals (aged 7-12 weeks) revealed both mass reductions at the level of tissue and morphological changes at the cellular level in *PTPN23^H/H^* mice. Dissection of individual fat deposits – including inguinal (IAT), mesenteric (MAT), perigonadal (GAT), perirenal (RAT), retroperitoneal (RPAT), and subscapular (SSAT) adipose tissues - demonstrated reductions in mass, normalized to total body mass, for all white adipose tissues examined (Figure 3A and 3B). However, similar reductions in mass were not observed in brown adipose tissue (BAT) or in liver, heart, lung, or gastrocnemius (Figure 3B and 3C). Reduction in subdermal white adipose tissue was especially apparent through histological examination of the skin, where *PTPN23^H/H^* mice had drastically reduced adipose (Figure 3D). Additionally, histological examination of IAT in *HDPTP^h/h^* mice showed reduction in adipocyte volume (Figure 3E). Collectively, these patterns suggest that adipogenesis can occur when HD-PTP is lacking, but lipid storage within the adipocytes is perturbed. More generally these data indicate that loss of HD-PTP function in *PTPN23^H/H^* mice results in generalized lipodystrophy.

**Figure 3.**
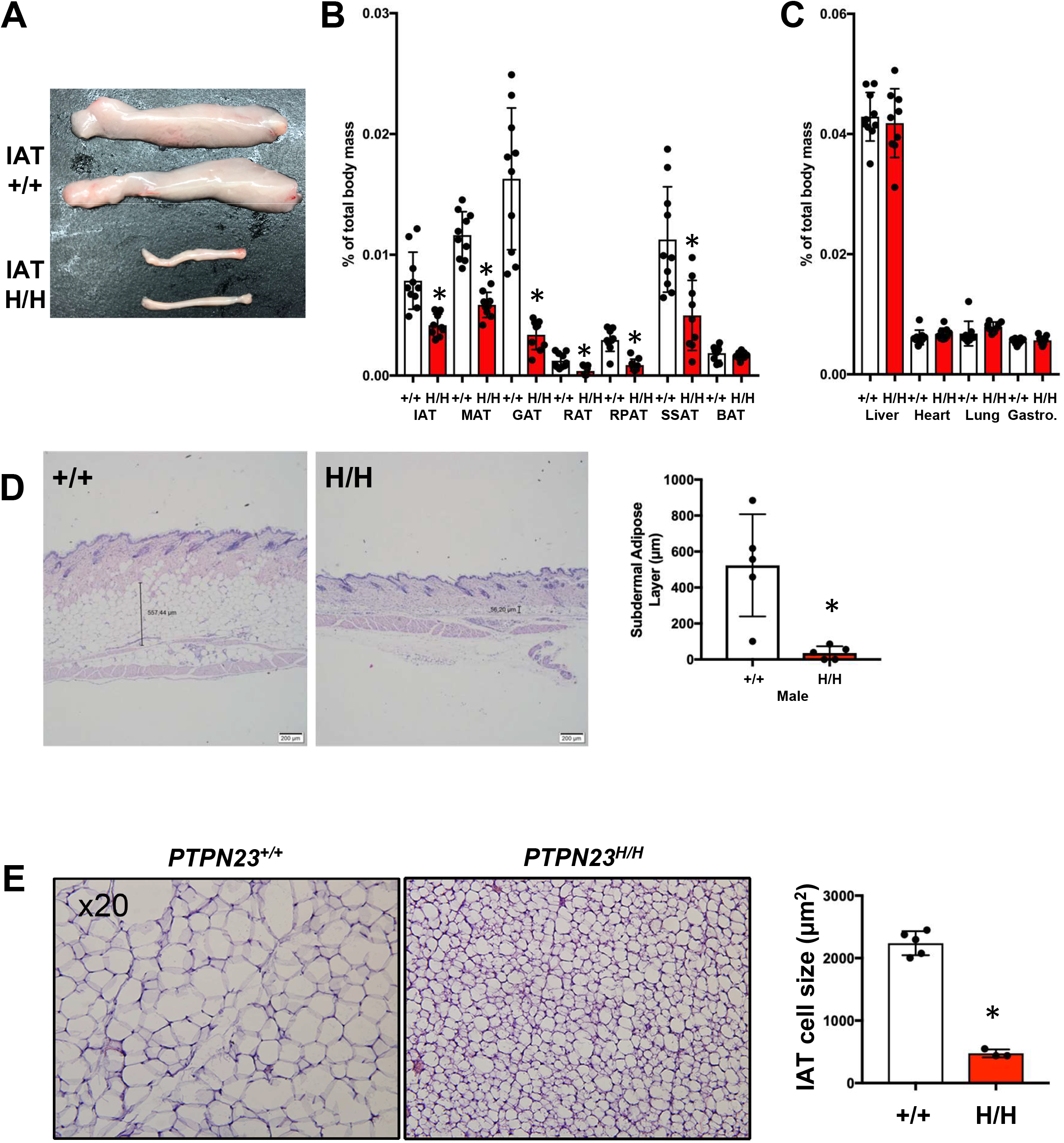
*PTPN23^H/H^* animals exhibit reduced white fat accumulation. **A.** Representative image of inguinal adipose tissue from adult *PTPN23^+/+^* and *PTPN23^H/H^* mice. **B,C.** Analysis of individual adipose tissues (**B**) and organs (**C**) from adult (age matched 6 week to 3 month old) male and female *PTPN23^+/+^* and *PTPN23^H/H^* mice normalized to total body mass. Adipose tissues examined: inguinal (IAT), mesenteric (MAT), gonadal (GAT), retroperitoneal (RPAT), subscapular (SSAT) and brown (BAT). Organs examined: liver, heart, lung, gastrocnemius. (**D**) Representative image of subdermal adipose tissue from adult *PTPN23^+/+^* and *PTPN23^H/H^* male mice. Quantitation of subdermal adipose layer depth (μm) presented. (**E**) Representative image of inguinal adipose tissue from adult *PTPN23^+/+^* and *PTPN23^H/H^* male mice. *PTPN23^H/H^* animals exhibit reduced IAT cell size (μm^2^). Quantitation of IAT cell size (μm^2^) presented for 5 *PTPN23^+/+^* and 3 *PTPN23^H/H^* male animals. * indicates two-tailed t-test p-value ≤ 0.05.

### Analyses of IAT reveals defects in signal transduction pathways

To address alterations in protein levels and phosphorylation states of signaling factors in white adipose tissue in *PTPN23^+/+^* and *PTPN23^H/H^* mice, IAT was used as a source for reverse phase protein array (RPPA) analysis given the relative ease of IAT isolation and the phenotypic differences in IAT between *PTPN23^+/+^* and *PTPN23^H/H^* adult mice (Figure 3A, 3B and 3E). HD-PTP expression in inguinal adipose tissue from *PTPN23^H/H^* mice was reduced ~67% compared to *PTPN23^+/+^* (Figure 4A). RPPA analysis of IAT from 9 adult age-matched male *PTPN23^+/+^* and *PTPN23^H/H^* animals revealed that 183/377 queried protein species displayed statistically significant alterations with 91 factors increased and 92 factors decreased in *PTPN23^H/H^* IAT (Figure 4B, Figure S4). While *PTPN23^H/H^* IAT adipocytes exhibit reduced volume and lipid droplet size, RPPA analysis revealed *PTPN23^H/H^* IAT has increased levels of lipid synthesis enzymes Acetyl Co-A Synthase, Acetyl-CoA Carboxylase and Fatty Acid Synthase (Figure 4C), suggesting that *PTPN23^H/H^* IAT is subject to altered regulation of lipid storage rather than defects in the lipid biogenesis machinery itself. In addition, *PTPN23^H/H^* IAT exhibited reduced Caveolin-1 levels (Figure 4C), which is interesting as Caveolin-1 has been implicated in regulating lipid droplet homeostasis [77, 78] and mutations in *CAV1* are linked to lipodystrophy [79–82]. Additionally, Caveolin-1 physically interacts with HD-PTP [83]. To gain further insights into the differences between *PTPN23^+/+^* and *PTPN23^H/H^* IAT, pathway analysis of RPPA results was undertaken (Figure 4D). This analysis revealed that *PTPN23^H/H^* IAT exhibits hallmarks of reduced activation of the Breast Reactive and Core Reactive pathways while the Hormone Receptor and Cell Cycle pathways were stimulated. In addition, decreased signaling through Receptor Tyrosine Kinase, RAS/MAPK, and PI3/AKT pathways was evident, including reduced EGFR activation and reductions in the levels and phosphorylation states of multiple components of the RAS/MAPK, and PI3/AKT pathways (Figure 4E). Consistent with this finding, western blotting revealed reduced activation of mTOR, Erk1/2 and Akt(S473) as well as increased levels of Insulin Receptor in *PTPN23^H/H^* IAT (Figure 4F–4I). As insulin is a driver of lipid storage in IAT [84], these results suggest that perturbed insulin signaling in IAT may contribute to reduced lipid storage in *PTPN23^H/H^* mice.

**Figure 4.**
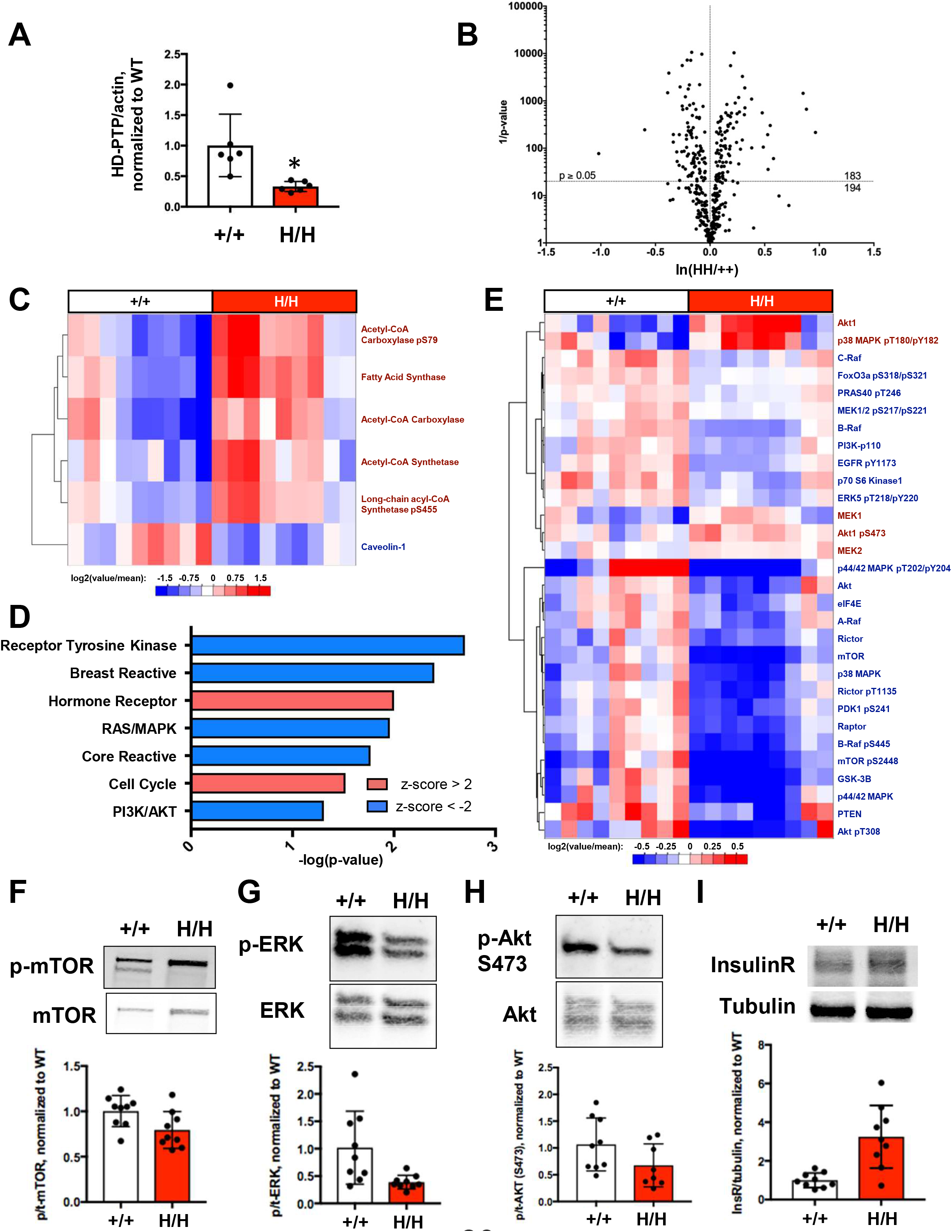
Adult *PTPN23^H/H^* IAT exhibit hallmarks of reduced receptor tyrosine kinase signaling. **A.** HD-PTP level, normalized to actin, in inguinal adipose tissue from adult *PTPN23^+/+^* and *PTPN23^H/H^* male animals. *PTPN23^H/H^* animals exhibit 67% reduction in HD-PTP level. * indicates two-tailed t-test p-value ≤ 0.05. **B.** Volcano plot of adult male IAT RPPA analysis [ln(HH/++) vs 1/p-value]. 183 protein species exhibit significant variation between *PTPN23^H/H^* and *PTPN23^+/+^* IAT. Dashed line indicates 0.05 p-value. **C.** Heat map of significantly altered (two-tailed t-test p-value ≤ 0.05) lipid metabolism factors and caveolin-1 in RPPA analysis. Labels for factors elevated in *PTPN23^H/H^* are red, while labels for factors reduced in *PTPN23^H/H^* are blue. **D.** Pathway analysis of male adult IAT RPPA results indicating significantly altered pathways, including Receptor Tyrosine Kinase (RTK), Ras/MAP Kinase, and PI3Kinase/Akt pathways (highlighted in green). The −log_10_ of the p-value is plotted. Activated pathways with z-score > 2 are pink, and inhibited pathways with z-score < −2 are blue. **E.** Heat map of some of the significantly altered (two-tailed t-test p-value ≤ 0.05) signaling components in *PTPN23^H/H^* adult male IAT as determined by RPPA. Labels for factors elevated in *PTPN23^H/H^* are red, and labels for factors reduced in *PTPN23^H/H^* are blue. **F-H**. Representative immunoblots and quantitation analyzing activation of mTOR (**F**), ERK (**G**) and Akt (**H**) in *PTPN23^+/+^* and *PTPN23^H/H^* adult male IAT. **I.** Representative immunoblot and quantitation analyzing level of Insulin Receptor β subunit, normalized to tubulin, in *PTPN23^+/+^*and *PTPN23^H/H^* adult IAT. * indicates two-tailed t-test p-value ≤ 0.05.

### Altered insulin signaling in adult IAT is subject to extrinsic and intrinsic factors

Altered insulin signaling could result from cell intrinsic or extrinsic changes in *PTPN23^H/H^* mice. We first examined circulating insulin and glucose levels as extrinsic hallmarks of altered insulin signaling. Circulating insulin was reduced in both male and female *PTPN23^H/H^* mice (Figure 5A), however blood glucose levels were indistinguishable from *PTPN23^+/+^* under freely-fed and fasting conditions (Figure 5B and 5C). These observations suggest that *PTPN23^H/H^* mice maintain normal glycemic index through secreting less insulin than *PTPN23^+/+^* mice, implicating insulin hypersensitivity rather than the insulin resistance suggested by RPPA analysis of IAT. To test this model, glucose and insulin tolerance tests were performed. Glucose tolerance tests revealed no difference between *PTPN23^+/+^* and *PTPN23^H/H^* mice in circulating glucose levels in response to a glucose bolus (Figure 5D), indicating *PTPN23^H/H^* mice do not exhibit insulin resistance. In contrast, insulin tolerance tests revealed that *PTPN23^H/H^* mice are hypersensitive to exogenous insulin (Figure 5E). This insulin hypersensitivity and reduced circulating insulin under non-fasting conditions suggest that cell extrinsic factors are one contributor to reduced insulin signaling in *PTPN23^H/H^* IAT. Insulin-induced signaling was examined *in vivo* to complement the insulin tolerance test. Fasting mice were stimulated with insulin, and induced signaling was examined in IAT as well as target tissues of insulin, including liver and skeletal muscle, shortly after injection. Whereas *PTPN23^+/+^* and *PTPN23^H/H^* IAT and liver exhibit comparable Akt activation at this early time point (Figure 5F and 5G), *PTPN23^H/H^* skeletal muscle exhibits enhanced Akt activation (Figure 5H). IAT did not exhibit defects in insulin-induced AKT activation at 10 minutes *in vivo* in contrast to reduced AKT activation in freely fed mice, therefore we performed *in vitro* insulin stimulation of IAT to evaluate signaling over a timecourse rather than a single timepoint. While basal levels and initial stimulation of phospho-AKT were comparable, levels of insulin-induced Akt activation at extended timepoints were reduced in *PTPN23^H/H^* IAT (Figure 5I) despite elevated levels of Insulin Receptor (Figures 4I and 5J). These results indicate that HD-PTP deficiency impacts insulin signaling distinctly in different tissues, conferring insulin resistance or sensitivity depending on the context. In total, these observations highlight both intrinsic and extrinsic contributions to reduced insulin signaling in IAT in adult HD-PTP deficient mice.

**Figure 5.**
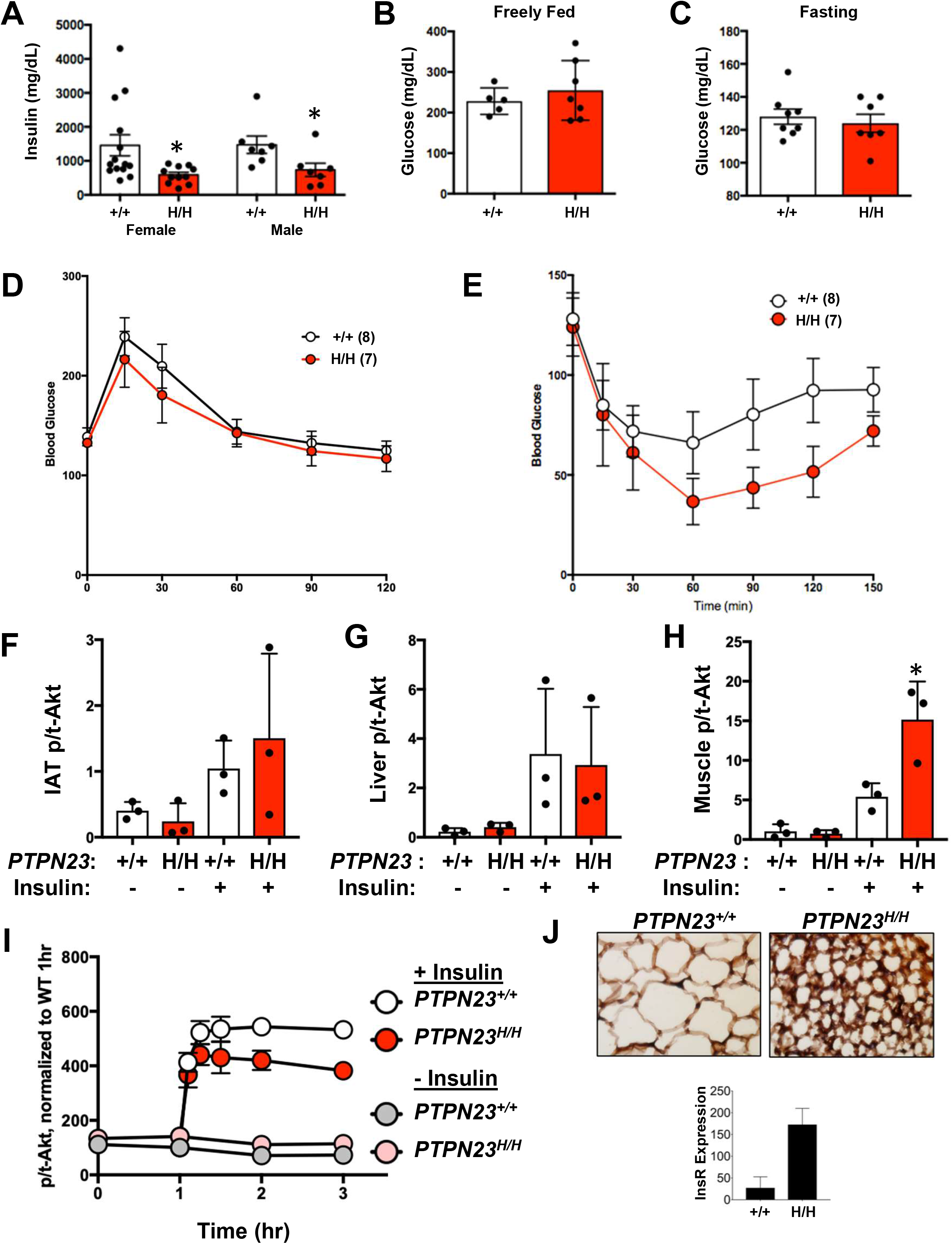
*PTPN23^H/H^* adult animals exhibit insulin hypersensitivity but reduced insulin signaling in IAT *in vitro*. **A.** Circulating insulin level (mg/dL) in *PTPN23^+/+^* and *PTPN23^H/H^* female and male freely-fed adult animals. **B,C.** Blood glucose level in freely-fed (**B**) or 6 hour starved (**C**) *PTPN23^+/+^* and *PTPN23^H/H^* adult female animals. **D.** Glucose tolerance test in which 1.25g/kg glucose was administered intraperitoneally to *PTPN23^+/+^* and *PTPN23^H/H^* adult female animals, and circulating glucose levels were assessed 15, 30, 60, 90 and 120 minutes later. Glucose levels are comparable between *PTPN23^+/+^* and *PTPN23^H/H^* animals under these conditions. **E.** Insulin tolerance test in which 0.5mU/kg insulin was administered intraperitoneally to *PTPN23^+/+^* and *PTPN23^H/H^* adult female animals, and circulating glucose levels were assessed 15, 30, 60, 90, 120 and 150 minutes later. *PTPN23^H/H^* animals exhibit reduced glucose levels under these conditions. **F-H.** Starved *PTPN23^+/+^* and *PTPN23^H/H^* adult animals were stimulated with 0.5mU/kg insulin administered intraperitoneally, and tissues – inguinal adipose tissue (**F**), liver (**G**) and muscle (**H**) – were harvested 10 minutes later. Akt activation was assessed by immunoblotting for phospho-Akt S473 and total Akt, and the ratio of phospho/total-Akt is presented. **I.** *In vitro* stimulation of *PTPN23^+/+^* and *PTPN23^H/H^* inguinal adipose tissue in which 100ng/ml insulin was administered to suspended IAT fragments (at 1 hr timepoint) and lysates were generated 5, 15, 30, 60 and 120 minutes after stimulation. Akt activation was assessed by immunoblotting for phospho-Akt S473 and total Akt, and the ratio of phospho/total-Akt normalized to +/+ prior to stimulation is presented. **J.** Representative image of Insulin Receptor β staining of adult *PTPN23^+/+^* and *PTPN23^H/H^* animal inguinal adipose tissue. * indicates two-tailed t-test p-value ≤ 0.05.

### HD-PTP deficiency perturbs IAT adipogenesis

The extensive changes between adult *PTPN23^+/+^* and *PTPN23^H/H^* mice evident in both RPPA analysis and characterization of the fat tissues led us to explore variation earlier in post-natal development that could represent the primary driver of the phenotypes observed in the adult animals. Differences in mass between *PTPN23^H/H^* and *PTPN23^+/+^* animals were observed at both 10 and 21 days (Figure 6A), and reduced body fat composition in *PTPN23^H/H^* animals was also apparent at these early time points (Figure 6B). However, the difference between *PTPN23^H/H^* and *PTPN23^+/+^* was less severe than in >6-week-old animals (Figure 2C and 2D). Consistent with this observation, HD-PTP expression in 14-day IAT is reduced only ~50% (Figure 6C) compared to the ~67% reduction observed in adult IAT (Figure 4A). RPPA analysis of IAT isolated from 14-day-old *PTPN23^+/+^* and *PTPN23^H/H^* animals (n=3 female and 3 male for each) identified 32/377 queried protein species displaying statistically significant alterations (Figure 6D, Figure S6). IAT from 14-day-old *PTPN23^H/H^* mice exhibited increased levels of Insulin Receptor and IGF1R as well as increases in IRS2, mTOR and p70 S6 Kinase 1; however, InsR/IGF1R phosphorylation and downstream activation of Ribosome S6 were reduced, suggesting reduced insulin signaling in the 14-day *PTPN23^H/H^* IAT similar to adult IAT (Figure 6E). In contrast to the adult IAT, 14-day-old *PTPN23^H/H^* IAT exhibited decreased levels of lipid synthesis enzymes Acetyl Co-A Synthase, Acetyl-CoA Carboxylase and Fatty Acid

**Figure 6.**
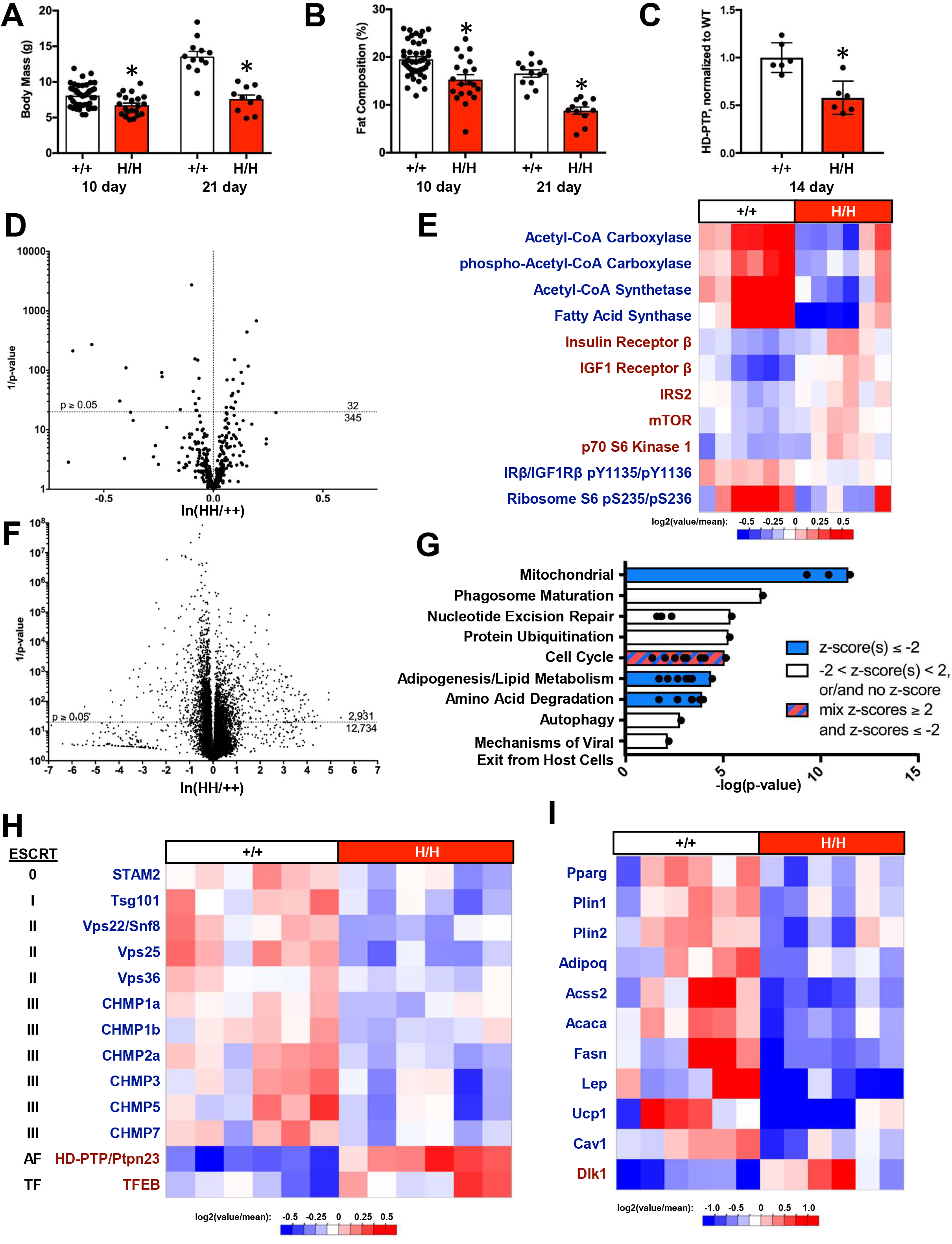
*PTPN23^H/H^* 14-day old animals exhibit exhibit hallmarks of reduced adipogenesis and reduced Insulin signaling. **A,B.** Total body mass (**A**) and total body fat composition analysis (**B**) of 10 and 21 day old *PTPN23^+/+^* and *PTPN23^H/H^* animals. **C.** HD-PTP level in inguinal adipose tissue from 14 day old *PTPN23^+/+^* and *PTPN23^H/H^* animals. *PTPN23^H/H^* animals exhibit 42% reduction in HD-PTP level. **D.** Volcano plot of 14-day old animal IAT RPPA analysis [ln(HH/++) vs 1/p-value]. 32 protein species exhibit significant variation between *PTPN23^H/H^* and *PTPN23^+/+^* IAT. Dashed line indicates 0.05 p-value. **E.** Heat map of significantly altered (two-tailed t-test p-value ≤ 0.05) lipid metabolism factors and some of the signaling factors in 14 day old animal IAT RPPA analysis. Labels for factors elevated in *PTPN23^H/H^* are red, while labels for factors reduced in *PTPN23^H/H^* are blue. **F.** Volcano plot of 14-day old animal IAT RNASeq analysis [ln(HH/++) vs 1/p-value]. 2,931 RNA species exhibit significant variation between *PTPN23^H/H^* and *PTPN23^+/+^* IAT. Dashed line indicates 0.05 p-value. **G.** Ingenuity canonical pathway analysis of transcripts significantly altered (p value ≤ 0.05) between 14-day old animal *PTPN23^H/H^* and *PTPN23^+/+^* IAT identified a number of pathways that clustered under the areas of Mitochondrial function, Phagososme maturation, Nucleotide excision repair, Protein ubiqutination, Cell cycle regulation, Adipogenesis/lipid metabolism, Amino acid degradation, Autophagy, and Mechanisms of viral exit from host cell. Bars indicate largest −log_10_ of the p-value within the cluster. Clusters with pathways with z-scores ≤ −2, suggesting inhibition, are colored blue. The Cell cycle regulation cluster is colored both pink and blue to indicate it contains pathways with z-scores ≤ −2 and z-scores ≥ 2. Clusters containing pathways without z-scores or with z-scores between −2 and 2 are colored white. **H.** Heat map of significantly altered (p value ≤ 0.05) ESCRT subunits and associated factors identified by RNASeq analysis of 14 day old animal IAT. **I.** Heat map of adipogenesis markers, Cav1 (encoding caveolin-1) and a preadipocyte marker (Dlk1) identified by RNASeq analysis as significantly altered (p value ≤ 0.05) between 14-day old animal *PTPN23^H/H^* and *PTPN23^+/+^* IAT.

Synthase and decreased phosphorylation of Acetyl-CoA Carboxylase compared to *PTPN23^+/+^* (Figure 6E). To gain additional information pertaining to these differences, RNASeq analysis was performed on IAT from 14-day-old *PTPN23^+/+^* and *PTPN23^H/H^* mice (n=3 female and 3 male animals for each). We identified 831 significantly up-regulated and 1,356 significantly down-regulated genes in *PTPN23^H/H^* 14-day IAT (Figure 6F). Ingenuity Pathway Analysis (IPA) of these altered genes identified a number of canonical pathways that may be relevant to altered IAT and HD-PTP function (Figure 6G). While the connections between HD-PTP and mitochondrial function, nucleotide excision repair, cell cycle and amino acid degradation pathways are unclear, the ESCRT machinery has been implicated in three of these identified canonical pathways: Protein Ubiquitination, Autophagy, and Mechanisms of Viral Exit from Host Cells. Further inspection revealed that 11 ESCRT subunits exhibited reductions in *PTPN23^H/H^* IAT (Figure 6H). IPA also identified alterations in the adipogenesis pathway and five additional pathways related to lipid homeostasis. Nine markers of adipogenesis exhibited reduced expression in *PTPN23^H/H^* IAT while Dlk1, a marker of preadipocytes, was elevated (Figure 6I). These results indicating reduced adipogenesis are consistent with reduced expression of Fatty Acid Synthase, Acetyl-CoA Synthase and Acetyl CoA Carboxylase evident in 14-day-old IAT RPPA analysis (Figure 6D). RPPA analysis indicated these factors are elevated in adult *PTPN23^H/H^* IAT (Figure 4C), consistent with the conclusion that the reduction in adipogenesis markers at 14 days of age is an indicator of reduced or delayed adipogenesis. IPA also identified several upstream regulators with altered activation state including Insulin Receptor (p-value 1.28e-4; z-score −3.56) and insulin (p-value 1.41e-4; z-score −2.391); these results are consistent with the reduced activation of InsR/IGF1R and Ribosome S6 apparent in RPPA analysis (Figure 6E). These results suggest that insulin signaling, a driver of adipogenesis, and the adipogenesis pathway are negatively impacted in developing IAT when HD-PTP levels are reduced.

These observations led to examination of HD-PTP’s role in adipogenesis and insulin signaling *in vitro*. To avoid *in vivo* extrinsic contributions to HD-PTP deficient preadipocyte behaviors, we utilized *PTPN23^-/-^* preadipocytes derived from *PTPN23^flox/flox^* animals. Preadipocytes were isolated from *PTPN23^flox/flox^* IAT and cultured *in vitro*, followed by *PTPN23* knockout via doxycycline-induced Cre expression and adipogenesis induction. Immunoblotting for markers of adipogenesis indicated reduced adipogenesis in *PTPN23^-/-^* preadipocytes, and this reduction was not observed in *PTPN23^flox/flox^* preadipoyctes (Figure 7A and 7B). These results are consistent with the reduced adipogenesis observed through RNASeq analysis of *PTPN23^H/H^* IAT. We next examined insulin signaling in *PTPN23* knockout preadipocytes. Insulin stimulation of *PTPN23^-/-^* preadipocytes resulted in reduced Akt activation compared to *PTPN23^+/+^* (Figure 7C), consistent with RPPA and RNASeq analysis of 14-day *PTPN23^H/H^* IAT and analysis of signaling in adult *PTPN23^H/H^* IAT. Taken together, these observations indicate that HD-PTP deficiency perturbs insulin signaling and adipogenesis in preadipocytes.

**Figure 7.**
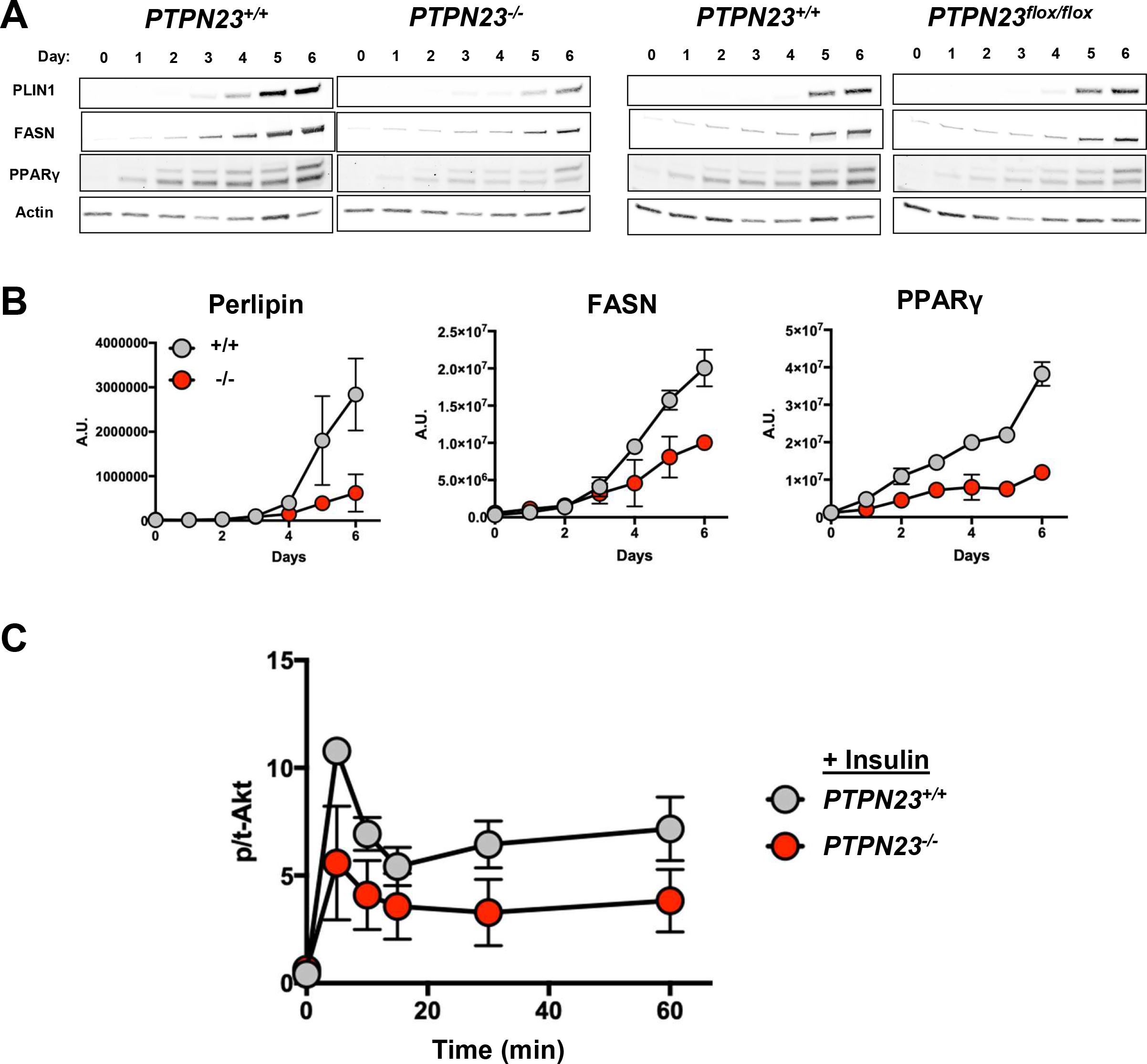
*PTPN23* knockout preadipocytes exhibit defects in adipogenesis and insulin signaling in vitro. **A, B.** *PTPN23^-/-^* preadipocytes - preadipocytes isolated from *PTPN23^flox/flox^* animals and knockout implemented *in vitro* by doxycycline-induced CRE expression subsequent to establishing culture – were subjected to *in vitro* adipogenesis procedure and lysates were generated for each of 7 days of differentiation. Adipogenesis of *PTPN23^+/+^* and *PTPN23^flox/flox^* preadipocytes without doxycycline exposure is presented in right panels. Adipogenesis markers Perilipin1 (PLIN1), Fatty Acid Synthase (FASN), and Peroxisome Proliferator-activated Receptor γ (PPARγ) were evaluated by immunoblotting (**A**). Actin was evaluated as a loading control. Quantitation of Perlipin, FASN and PPARγ expression in *PTPN23^+/+^* and *PTPN23^-/-^* cells is presented in **B. C**. *In vitro* stimulation of *PTPN23^+/+^* and *PTPN23^-/-^* adipocytes in which 100ng/ml insulin was administered and lysates were generated 5, 10, 15, 30, and 60 minutes after stimulation. Akt activation was assessed by immunoblotting for phospho-Akt S473 and total Akt, and the ratio of phospho/total-Akt is presented.

## Discussion

The ESCRT machinery functions together with associated Bro1 Domain Family members, including HD-PTP and ALIX, to execute membrane remodeling processes that are more diverse than initially appreciated. In addition to driving MVB biogenesis for receptor downregulation, the ESCRTs facilitate abscission during cytokinesis, the biogenesis of extracellular vesicles and many viruses, closure of the autophagosome, repairing plasma membrane lesions, and maintaining nuclear envelope integrity. The critical contributions of ESCRTs to mammalian physiology is highlighted by the embryonic lethality of mouse ESCRT knockout models [9–11] and the disease-associated mutations linked to ESCRT genes [13–25]. For example, the *PTPN23* (HD-PTP) mouse knockout model is embryonic lethal [12], and mutations in HDPTP/PTPN23 are linked to NEDBASS [26–30]. To circumvent embryonic lethality of *PTPN23* knockout in mice and gain further insights into HD-PTP-mediated processes contributing to mammalian physiology, we developed a PTPN23 hypomorphic allele. *PTPN23^H/H^* animals develop to adulthood but exhibit striking phenotypes of reduced survival and lipodystrophy. Our analyses support the conclusion that HD-PTP function is required to maintain homeostasis in the adult mouse in addition to its requirement in embryonic development [12] and its role as a tumor suppressor [71]. We did not observe increased tumorigenesis in the *PTPN23^H/H^* animals, as predicted from studies with *PTPN23^-/+^* animals [71]. We presume that shortened *PTPN23^H/H^* life span has occluded this phenotype. The most striking phenotype of *PTPN23^H/H^* animals is their reduced size, largely attributable to the reduction in fat composition. Decreased fat composition may explain why increased total energy expenditure was observed in *PTPN23^H/H^* animals as lean tissue is more metabolically active than fat tissue; however, at present it is not possible to eliminate altered metabolism due to HD-PTP deficiency as contributing to altered body composition.

Characterization of *PTPN23^H/H^* IAT revealed an unexpected connection among the ESCRTs at the transcriptional level. RNASeq analysis revealed reduction in components of ESCRT-0 (STAM2), ESCRT-I (Tsg101), ESCRT-II (Vps22/Snf8, Vps25, Vps36), and ESCRT-III (CHMP1a, CHMP1b, CHMP2a, CHMP3, CHMP5, CHMP7) in *PTPN23^H/H^* IAT. In this context where reduction of HD-PTP has compromised ESCRT functions such as MVB biogenesis, the cell appears to have downregulated other machinery contributing to this process. A second alternative is that reduced growth factor signaling in *PTPN23^H/H^* IAT is contributing to this effect, implicating growth factor signaling in upregulating ESCRTs at the transcriptional level in normal cellular contexts as a potential feedback mechanism. As reduced ESCRT function has been linked with tumorigenesis and a number of congenital conditions, further exploration of the basis of coordinated ESCRT expression may provide a novel means to alter flux (e.g. enhance MVB biogenesis) for therapeutic benefit.

A prominent and unexpected phenotype of our *PTPN23^H/H^* mice was lipodystrophy, and we sought to understand how HD-PTP mechanistically contributes to fat accumulation. Our results indicate that both cell extrinsic and cell intrinsic defects contribute to lipodystrophy. HD-PTP deficiency results in systemic insulin hypersensitivity, as indicated by insulin tolerance tests and examination of insulin induced signaling in skeletal muscle. We infer that observed reductions in circulating insulin in the *PTPN23^H/H^* animals is a response to maintain normal glycemic levels, and that reduced circulating insulin contributes to reduced lipid storage in adipose tissue - consistent with reduced signaling evident by RPPA pathway analysis of freely-fed adult *PTPN23^H/H^* mice. Inherent within this model is the idea that adipose tissues do not exhibit the insulin-hypersensitivity evident in skeletal muscle. This conclusion is supported by absence of enhanced insulin-induced *PTPN23^H/H^* IAT signaling *in vivo*, the reduction in insulin-induced *PTPN23^H/H^* IAT signaling *in vitro*, and the reduction in insulin-induced signaling in *PTPN23* knockout preadipocytes *in vitro*. These results suggest HD-PTP makes tissue-specific contributions to insulin signaling negatively impacting insulin signaling in muscle while positively impacting insulin-signaling in adipose tissue. Mutation of *CAV1* (encoding caveolin-1) is linked to Congenital Generalized Lipodystrophy Type 3 [85, 86], and HD-PTP interacts with caveolin-1 [83]. These observations raise the possibility that HD-PTP and caveolin may function together to promote insulin-signaling and lipid storage in adipose tissue. Alternatively, the reduction in caveolin-1 in *PTPN23^H/H^* IAT, apparent in both adult and 14-day IAT, may contribute to altered insulin signaling rather than HD-PTP and caveolin-1 functioning together. While HD-PTP expression is similar in both adipose tissue and skeletal muscle, caveolin-1 is most strongly expressed in adipocytes [87, 88]. This difference suggests that while caveolin-1 and HD-PTP may promote signaling in adipose tissue, caveolin-1-independent HD-PTP function may inhibit insulin signaling in muscle. Inhibition of signaling in skeletal muscle is consistent with the classical mode of ESCRT-regulated receptor downregulation via MVB sorting. The potentiation of signaling in adipose tissue by ESCRTs is less understood but has been observed with Toll signaling in fruit flies [72]. Further understanding how HD-PTP, the ESCRTs and caveolin-1 promote insulin and Toll signaling rather than negatively regulating these events will be important to pursue, and this *PTPN23* hypomorphic system provides a useful model for dissecting this process.

## Materials and Methods

### Mouse strains

Mouse protocols were reviewed and approved by the Mayo Clinic Institutional Animal Care and Use Committee (IACUC). *PTPN23* hypomorphic allele (H) containing a neomycin selectable cassette in the reverse orientation between exons 6 and 7 was generated by standard gene targeting techniques and embryonic stem cell technology. *PTPN23^H/+^* mutant mice were maintained on a mixed 129Sv/E x C57BL/6 genetic background or backcrossed 6 generations to the C57BL/6 strain, and *PTPN23^H/+^* intercrosses were used to generate *PTPN23^H/H^* mice. *PTPN23^H/H^* mice in the mixed background or C57BL/6 background behaved equivalently. The neomycin cassette was removed by crossing *PTPN23^H/+^* mice with FLP-expressing mice to generate *PTPN23^flox^*. Intercrosses with R26-M2rtTA and TetOp-CRE mice were used to generate *PTPN23^flox/flox^ R26-M2rtTA/R26-M2rtTA +/+* and *PTPN23^flox/flox^ R26-M2rtTA/R26-M2rtTA +/TetOp-CRE* mice for intercrossing to generate *PTPN23^flox/flox^ R26-M2rtTA/R26-M2rtTA +/TetOp-CRE* mice used to isolate preadipocytes for doxycycline-induced knockout studies. Exons 5 and 6 were removed by crossing *PTPN23^H/+^* mice with *Hprt-Cre* transgenic mice to generate the null allele (-). *PTPN23^-/+^* intercrosses were performed to examine the viability of *PTPN23^-/-^* mice (Figure 1B). *PTPN23^H/-^, PTPN23^-/+^, PTPN23^H/+^* and *PTPN23^+/+^* pups were generated by crossing *PTPN23^H/+^* and *PTPN23^-/+^* mice; *PTPN23^H/H^* and *PTPN23^+/+^* pups were generate by intercrossing *PTPN23^H/+^* mice. Male and female mice were used for experimentation.

### Body composition, physiological and behavioral analyses

Body composition and total body fat were determined using an EchoMRI-100 QNMR instrument (Echo Medical Systems) as previously described [89]. Food intake, activity and metabolic rate were measured using a Comprehensive Lab Animal Monitoring System (CLAMS) as described in [90]. Individual experiments analyzed ~6-week-old *PTPN23^H/H^* and *PTPN23^+/+^* mice (n=4 female and male for each) and were repeated for total of 3 sets of animals. Energy expenditure was normalized to either total body mass or lean body mass, as determined using the EchoMRI-100. Individual fat depot mass and additional organs (liver, heart, lung, gastrocnemius) were determined through dissection and normalized to total body mass. Skin and fat (IAT) were fixed in 10% normal buffered formalin, embedded in paraffin, and sections were stained with hematoxylin and eosin (H&E) as previously described [89].

### RNA isolation, library preparation, sequencing and bioinformatics analyses

RNA was extracted from cryo-pulverized inguinal adipose tissue from freely-fed 14-day-old mice (n=3 male and female for *PTPN23^H/H^* and *PTPN23^+/+^*) using the RNeasy Mini Kit (74104; Qiagen) according to manufacturer’s instructions. The on-column DNase digestion step was included. RNA library preparation, sequencing and downstream analysis were performed as previously described [91, 92]. About 200 ng RNA was used for RNA library preparation. Sequencing was performed by the Mayo Clinic Molecular Biology Core Facility. Genes with significantly altered expression between *PTPN23^H/H^* and *PTPN23^+/+^* IAT were analyzed with Ingenuity Pathway Analysis (Qiagen) to identify impacted pathways and candidate upstream drivers. Expression variation between male and female animals was also examined to confirm results. Heat maps were created with the Next-Generation Clustered Heat Map (NG-CHM) Viewer web interface [93].

### Western Blot and Reverse Phase Protein Array (RPPA) analyses

Tissues were flash frozen, cryo-pulverized, and resuspended in Reverse Phase Protein Array (RPPA) Lysis Buffer [1% Triton X-100, 50mM HEPES, pH 7.4, 150mM NaCl, 1.5mM MgCl2, 1mM EGTA, 100mM NaF, 10mM Na Pyrophosphate, 1mM Na3VO4, 10% glycerol, 100μm AEBSF (Fisher BP2644-100), with Complete EDTA-free Protease Inhibitor Cocktail Tablet (Roche 04693132001) and PhosSTOP Phosphatase Inhibitor Cocktail Tablet (Roche 04906837001)]. Cleared lysate concentrations were measured with Bio-Rad Bradford protein assay and normalization was performed. Cleared lysates were combined with 5x Laemmli Sample Buffer, denatured, resolved by SDS-PAGE, and transferred to nitrocellulose. Western blotting with the following primary antibodies was performed: anti-HD-PTP V domain rabbit polyclonal antibody (1:500, provided by Robert Piper, University of Iowa), anti-HD-PTP F-4 mouse monoclonal antibody (1:100, sc-398711, Santa Cruz Biotechnology), anti-actin rabbit polyclonal antibody (1:1000, A2066, Sigma Aldrich), anti-β-actin mouse monoclonal antibody (1:10000, A5441, Sigma Aldrich), anti-α-tubulin mouse monoclonal antibody (1:1000, T6199, Sigma Aldrich), p-mTOR S2448 D9C2 rabbit monoclonal antibody (1:1000, 5536, Cell Signaling), total mTOR L27D4 mouse monoclonal antibody (1:1000, 4517, Cell Signaling), p-p42/44 MAPK (ERK1/2) Thr202/Tyr204 D13.14.4E rabbit monoclonal antibody (1:2000, 4370, Cell Signaling), total-p42/44 MAPK (ERK1/2) 3A7 mouse monoclonal antibody (1:2000, 9107, Cell Signaling), p-AKT-S473 D9E rabbit monoclonal antibody (1:1000, 4060, Cell Signaling), total Akt 40D4 mouse monoclonal antibody (1:1000, 2920, Cell Signaling), Insulin Receptor β CT-3 mouse monoclonal antibody (1:1000, AHR0271, ThermoFisher), Perilipin-1/PLIN1 D1D8 rabbit monoclonal antibody (1:1000, 9349, Cell Signaling), FASN C20G5 rabbit monoclonal antibody (1:1000, 3180, Cell Signaling), and PPARγ C26H12 rabbit monoclonal antibody (1:1000, 2435, Cell Signaling). Blots were developed with fluorescent secondary antibodies – goat anti-rabbit IRDye 680LT (1:5000, 926-68021, Li-Cor), goat anti-mouse IRDye 800CW (1:5000, 926-32210, Li-Cor). Signal was detected using the Odyssey Infrared Imager with ImageStudio software (LiCor) and analyzed using ImageJ (Schneider *et al*.,2012), ImageQuantTL (GE Life Sciences) and Prism 7 (GraphPad) to calculate the ratio of protein/loading control (actin or tubulin) or phospho/total protein. RPPA was performed by the MD Anderson Functional Proteomics RPPA Core Facility with 377 antibodies as described in [94]. Heat maps were created with the Next-Generation Clustered Heat Map (NG-CHM) Viewer web interface [93].

### Insulin assessment, glucose and insulin tolerance tests, insulin stimulation, and adipocyte differentiation

Serum Insulin levels were quantified using the Mouse Metabolic Discovery Array (MRDMET12, Eve Biotechnologies). Glucose tolerance test and insulin tolerance tests were performed as previously described [95]. Preadipocyte isolation was performed by mincing IAT in HBSS and digesting with Type 2 collagenase for ~1hr at 37 degrees. Preadipocytes were pelleted by 10min, 1200rpm spin at room temperature. Preadipocytes were plated in αMEM with 10% heat inactivated calf serum (CFS) and incubated overnight. The following day, preadipocytes were recovered with trypsin, replated at 50,000 cells/cm2 in αMEM 10% CFS, and cultured with 10% CO2, 3% O2 changing media every other day. Knockout of *PTPN23* was induced through addition of 1ug/ml doxycycline to the media of *PTPN23^flox/flox^ R26-M2rtTA/R26-M2rtTA +/TetOp-CRE* preadipocytes. Cells were cultured a minimum of 3 days before initiating adipogenesis. Preadipocyte differentiation was initiated at 85-90% confluence by feeding with Differentiation Medium (DMEM:F12 10% FBS, 1ug/ml insulin, 250nM dexamethasone, 0.5mM IBMX, 2.5uM Rosiglitazone) for 48 hours. Preadipocytes were then refed with DMEM:F12 10% FBS, 1ug/ml insulin for 7days harvesting samples each day. In vitro stimulation of IAT was performed with minced IAT in HBSS with protease and phosphatase inhibitors. Stimulation with 100ng/ml Insulin was performed after 1hr equilibration, and samples were lysed in RPPA Lysis Buffer with a dounce homogenizer 5, 15, 30, 60 and 120 minutes after stimulation. Western blotting was used to assess Akt activation. In vitro stimulation of preadipocytes was performed following 1hr serum starvation in DMEM:F12. Stimulation with 100ng/ml Insulin was performed, and samples were lysed in RPPA Lysis Buffer 5, 10, 15, 30, and 60 minutes after stimulation. Western blotting was used to assess Akt activation.

## Materials and Correspondence

Contact David Katzmann for materials. Contact Robert Piper for HD-PTP V domain polyclonal antibody.

## Acknowledgements

We are grateful for assistance provided by the Mark McNiven and Daniel Billadeau as well as the members of the McNiven, Billadeau, Kirkland and van Deursen labs at Mayo Clinic throughout this project.

## Figure Legends

**Figure 1, Supplementary Information.**
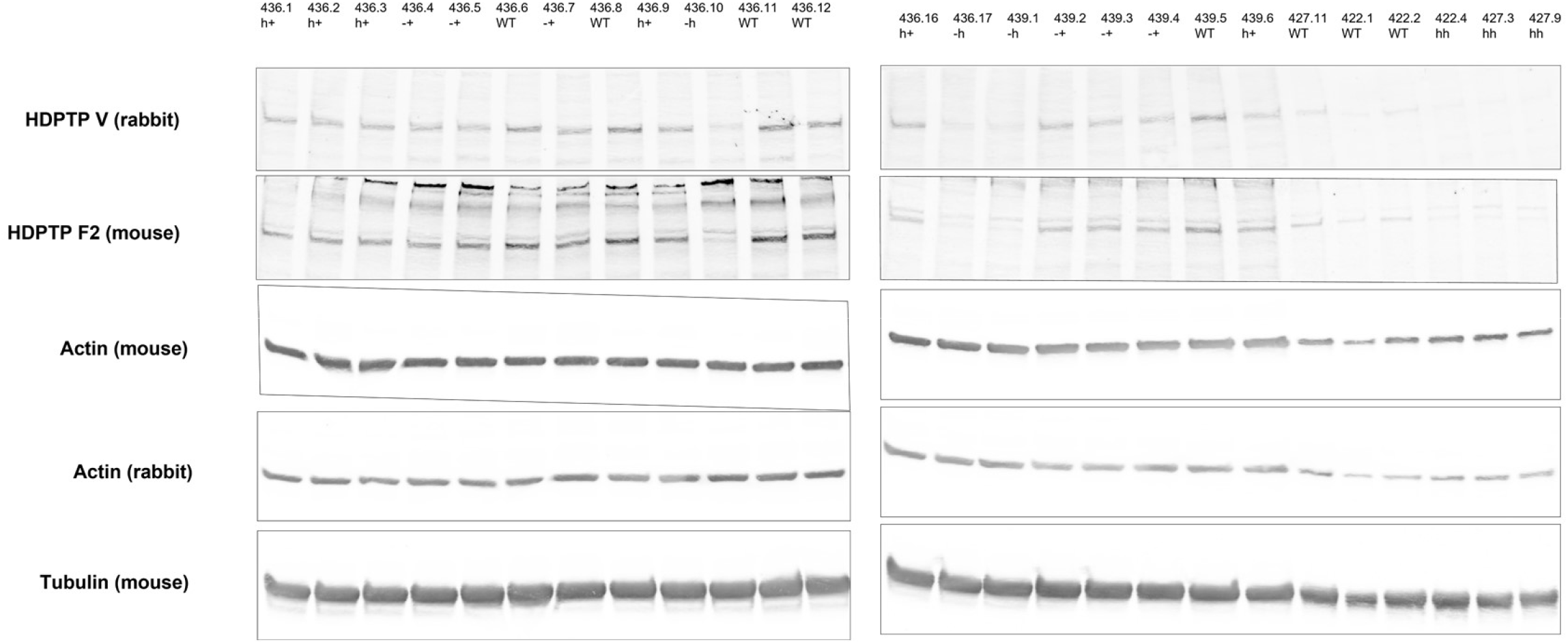
Analysis of HD-PTP expression in brain tissue from *PTPN23^+/+^, PTPN23^H/+^, PTPN23^-/+^, PTPN23^H/H^* and *PTPN23^H/-^* newborn pups. Western blotting was performed with 2 antibodies against HD-PTP, 2 antibodies against β-actin, and 1 antibody against tubulin. Quantitation of HD-PTP normalized to the actin and tubulin loading controls is presented in Figure 1C.

**Figure 2, Supplementary Information.**
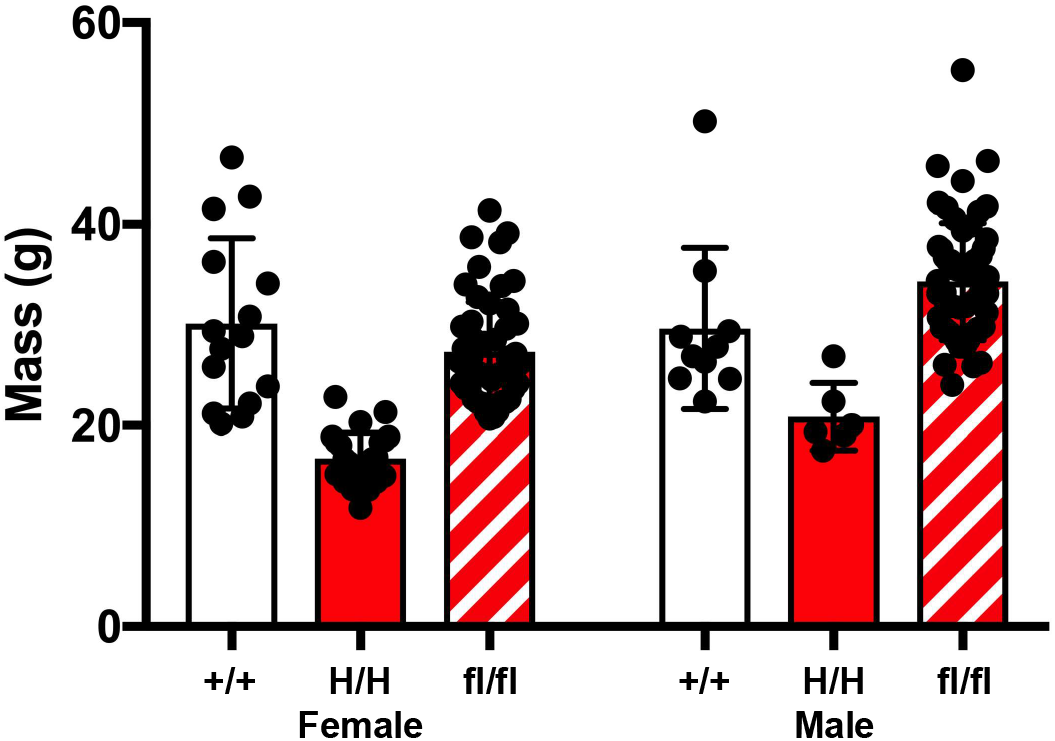
Total body mass of *PTPN23^+/+^, PTPN23^H/H^* and *PTPN23^flox/flox^* female and male adult animals.

**Figure 4, Supplementary Information.**
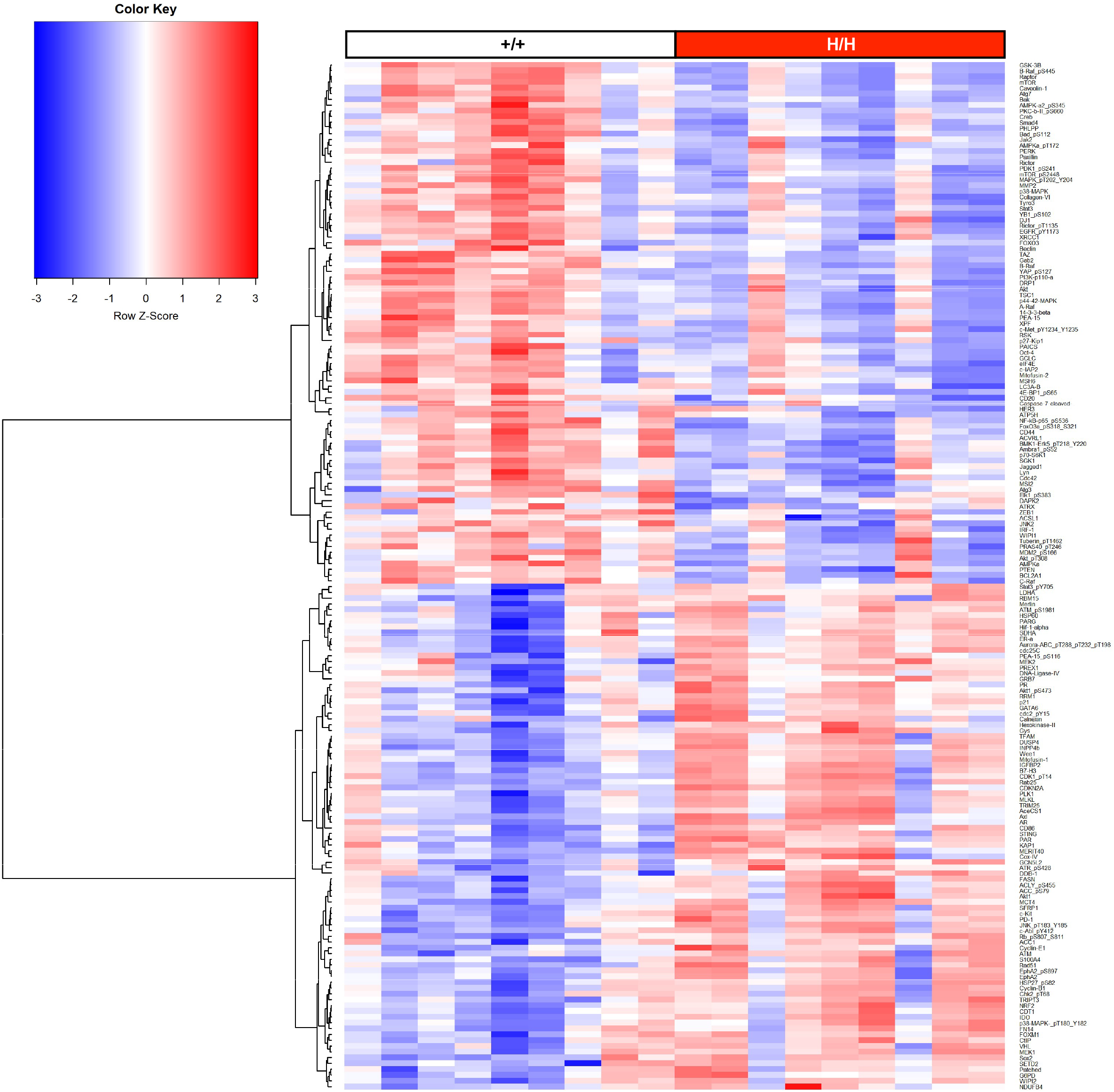
Heat map of 183 significantly altered (two-tailed t-test p-value ≤ 0.05) factors in RPPA analysis of adult male IAT.

**Figure 6, Supplementary Information.**
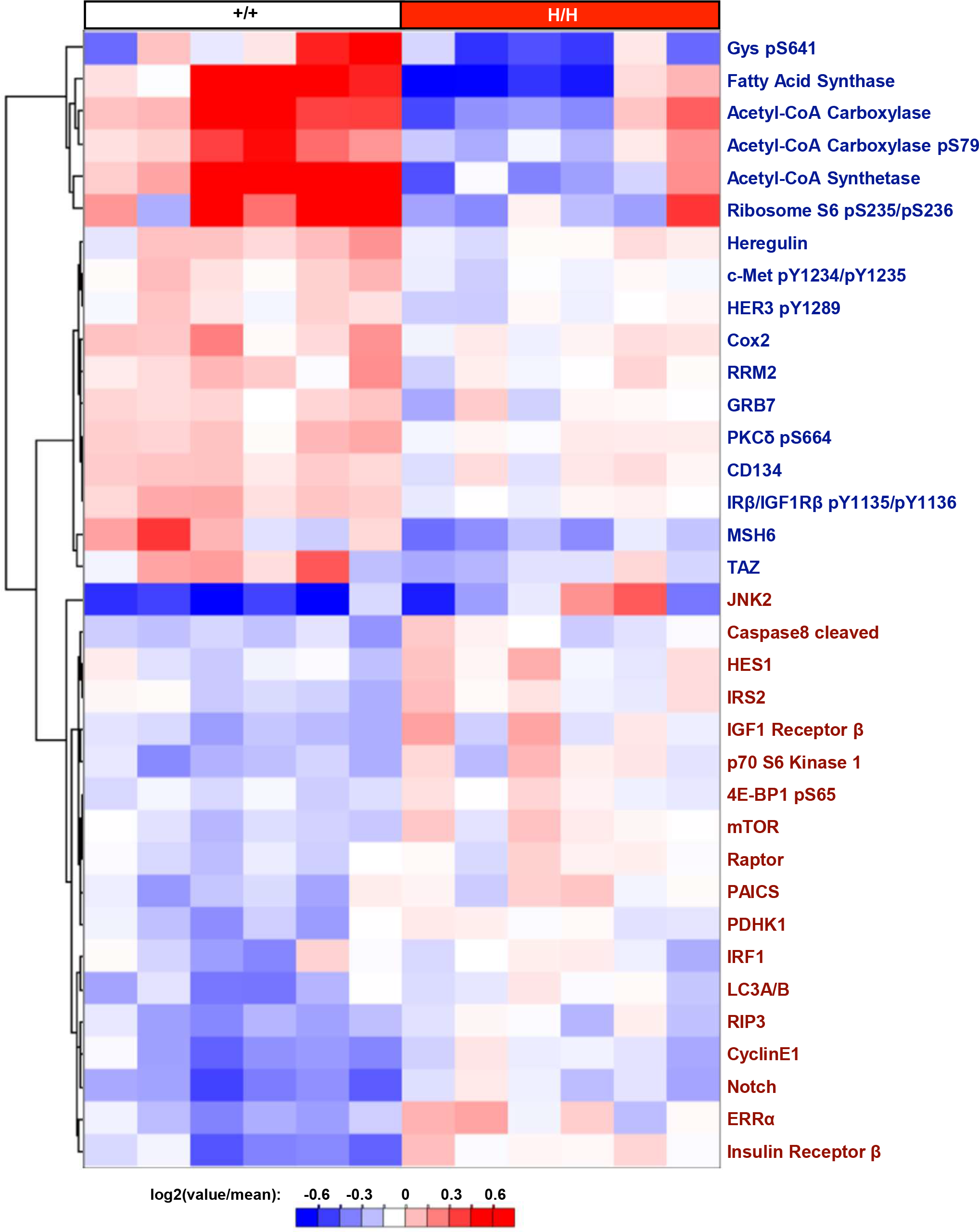
Heat map of all 32 significantly altered (two-tailed t-test p-value ≤ 0.05) factors in RPPA analysis of 14 day old animal IAT. Labels for factors elevated in *PTPN23^H/H^* are red, while labels for factors reduced in *PTPN23^H/H^* are blue.

